# Uniaxial tensile tests and Digital Image Correlation analysis for the mechanical characterization of human Fascia Lata under different decellularization treatments

**DOI:** 10.1101/2025.05.10.651796

**Authors:** Francesca Demontis, Giulia Loi, Edoardo Mazzotti, Mariana Rodriguez Reinoso, Federico Vecchio, Angelo Corso Faini, Agnese Leone, Elisa Camusso, Federico Genzano Besso, Cecilia Surace, Giuseppe Lacidogna, Domenico Scaramozzino

**Author notes:** These authors contributed equally (Francesca Demontis, Giulia Loi, Edoardo Mazzotti).

## Abstract

Fascia Lata (FL) is frequently employed as a graft source in reconstructive surgery. To minimize unwanted responses from the host immune system, several decellularization treatments have been proposed. Effective treatments should aim at avoiding the deterioration of the physical and mechanical properties of the implanted tissue. In this work, we carried out a mechanical characterization of FL specimens from human dead donors, both in their native-physiological condition and upon decellularization with three commonly used detergents, t-octyl-phenoxypolyethoxyethanol (Triton X-100), sodium dodecyl sulfate (SDS), and tri-n-butyl phosphate (TnBP). Uniaxial tensile tests were used to characterize the elastic stiffness and ultimate stresses of the tissue, and Digital Image Correlation (DIC) was applied to monitor the strain evolutions and meso-mechanical deformation responses. None of the investigated decellularization protocols was found to lead to a significant deterioration of the FL mechanical properties, suggesting the applicability of these chemical treatments for graft preparation and usage in the clinical practice. The application of DIC also allowed us to get a first estimate of the FL Poisson ratio as well as to draw the attention on the inhomogeneity of strain distributions, suggesting that the use of average engineering strains can lead to an oversimplification of the actual deformation field.

## 1. Introduction

The large amounts of surgical reconstructive procedures due to soft tissue traumatic injuries, acquired or iatrogenic defects, represent a significant health and economic burden. Generally, these procedures require the employment of autologous or allogenic grafts^1^. Fascia Lata (FL) has received a particular attention within the medical community since it represents a validated graft source and has been adopted in several types of reconstructive procedures^2^. FL grafts have been used as support in facial paralysis^3^ and facial reanimation^4^, for closure of enlarged tracheoesophageal punctures after laryngectomy-laryngopharyngectomy surgeries^5^, penile^6^ and pelvic^7^ reconstruction, ophthalmic plastic and reconstruction surgery^8^, reconstruction of full-thickness chest walls^9^, rhinoplasty interventions^10^, etc.

Biological graft materials are made of abundant extracellular matrix (ECM) and they generally need to undergo specific manufacturing methods that remove cellular and major histocompatibility complex components^11^. These methods include decellularization and chemical crosslinking to remove antigenic epitopes, DNA, and other biomolecules, while trying to minimize negative effects on the ECM features^12,13^. Decellularization methods can be classified into chemical-enzymatic or physical treatments, and encompass a wide variety of detergents, concentrations, and incubation times. The most common chemical-enzymatic decellularization methods make use of chemical detergents such as t-octyl-phenoxypolyethoxyethanol (Triton-X 100), sodium dodecyl sulfate (SDS), tri-n-butyl phosphate (TnBP), or enzymes such as deoxyribonucleases (DNAses) and trypsin^1^. The *in vivo* biodegradability of the ECM plays a key role in determining the host immune response and the remodeling processes inducing the clinical outcome^13,14^, thus constituting a driving factor for the choice of the optimal decellularization protocol. However, it is crucial that the implanted grafting material is also appropriate from a mechanical point of view^15^. Since the implanted graft is expected to withstand physiological loads, adequate mechanical strength and elasticity are necessary to allow mechanical loads to be correctly sustained by the surrounding tissues, while undergoing proper deformations and minimizing localized strains. Therefore, it is pivotal to properly characterize the mechanical behavior of the native tissue and how this is impacted by decellularization treatments.

Fascia Lata is an aponeurotic fascia, which encloses the tight muscles, and it is a dense and regular connective tissue with a multilayered organization, with collagen fibers disposed in each layer along specific directions. A comprehensive review of the mechanical characterization of human FL tissues can be found in a recent paper from Bonaldi et al^16^. Almost one century ago, Gratz et al. reported the first results arising from uniaxial tensile tests on human FL, assessing its ultimate strength and elasticity with different specimen arrangements^17,18^. Linear stiffness and elastic modulus of human samples under uniaxial tests were also investigated by Derwin et al.^19^, where the biomechanical properties of the tissue were studied in untreated, decellularized, and antibiotic-treated samples. Biaxial tests were performed by Eng et al.^20^ and Pancheri et al.^21^ to investigate the strength and stiffness of FL samples from adult goats, both in the longitudinal and transverse direction with respect to the orientation of collagen fibers. These tests confirmed that the material exhibits a strong anisotropic behavior, with higher stiffness in the longitudinal direction^20,21^. A comprehensive analysis of the mechanical behavior of human FL, also considering site specificity and gender differences, was carried out by Otsuka et al.^22^, who studied variabilities in stiffness, Young’s modulus, and hysteretic behavior in different samples. Pukšec et al.^23^ investigated the stress-strain characteristics of FL from fresh human dead donors, and compared them to those obtained from the muscle fascia and the dura mater. More recently, de Campos Azevedo compared the biomechanical properties of proximal and mid-thigh FL samples^24^, and Bonaldi et al.^25^ highlighted the strong anisotropic response with respect to the principal collagen fiber direction, also proposing a new constitutive model able to mimic intra- and interlayer interactions. All the studies reported above are characterized by large result variances^16^, which is mainly due to the: (1) intrinsic variability of the biological tissue properties; (2) different origin species (humans, goats, sheep) and specimen preparations; (3) different testing and experimental conditions, e.g., type of loading (uniaxial vs. biaxial, static vs. cyclic), speed of traction, gripping conditions to the loading device, etc.

Besides standard tensile tests, a few of the studies reported above also showed the use of complementary techniques which helped to interpret and integrate the tensile results, such as computer modeling (finite element method, analytical models) and full-field strain measurements. In particular, Digital Image Correlation (DIC) is an optical technique which allows to capture the displacements and strains of a surface under direct or indirect loadings. It is based on the application of a speckled pattern that is recognized by digital cameras and used for the assessment of the displacement field evolution through image-correlation algorithms^26^. This allows to investigate the full-field strain properties of the specimen, which is crucial in highly inhomogeneous materials such as biological samples. In recent years, DIC has found increasing application in biomechanics^27^, e.g., for the investigation of bovine cornea under constrained inflation conditions^28^, for the determination of strains in excised larynx vocal folds^29^, for the three-dimensional strain analysis of human tendons^30^, etc. A few years ago, Sednieva et al.^31^ made use of DIC together with motion analysis to assess the *in situ* strain evolution of FL during leg movement in three human dead donors. To the best of our knowledge, this was the only application of DIC to characterize deformation ranges of FL tissues, although not up to ultimate failure. In this work, we report the results of a mechanical investigation on human FL samples under uniaxial tension up to final collapse, complemented by DIC analysis, also exploring the impact that different chemical protocols have on the mechanical properties of the tissue.

## 2. Materials and Methods

### 2.1. Preparation of native FL specimens

Due to the unsteady availability of FL samples from human dead donors suitable for research purposes, mechanical tests were carried out in five separate experimental campaigns, i.e., in October 2020, January 2021, April 2021, February 2022, and May 2022. All FL tissues were carefully withdrawn from human dead donors in the operating room and placed for ∼10 minutes into a sterilized container filled with 10 liters of sanitizer at 5% concentration. After assessing that the tissue had not been contaminated before or during the removal, all samples were frozen at –80 °C, thawed the day before the mechanical tests, and then left at room temperature (Figure 1a). The tissues were then cut into rectangular samples (Figure 1b), so that the longer side followed the main direction of collagen fibers. We managed to collect a total of sixteen rectangular specimens from native FL, which were analyzed in the five experimental campaigns (see Table S1 in the Supplementary Material). The dimensions of the specimens were carefully measured via a digital caliber (for the width) and a digital micrometer (for the thickness). Several measurements on different points of the specimens were carried out, so that mean and standard deviation values of the geometrical dimensions were extracted. The cross-sectional areas of the samples, needed for stress calculations, were computed based on width and thickness values, applying the propagation of standard deviation errors (see Table S1). The initial length *L*_*0*_ of the sample was evaluated as the distance between the clamps in the testing machine at the beginning of the test (see Section 2.2).

**Figure 1.**
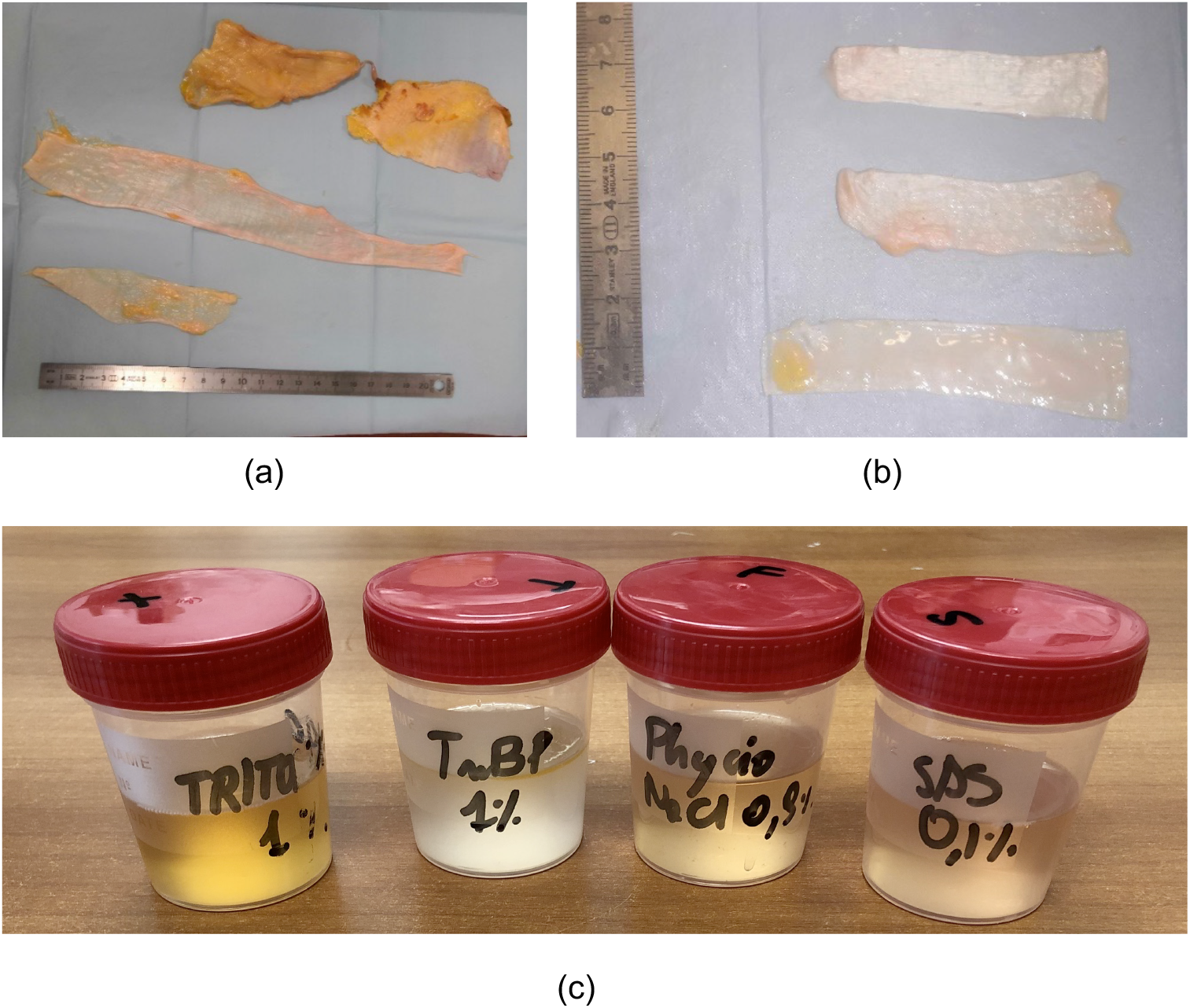
(a) Human Fascia Lata (FL) tissue from which individual specimens were collected; (b) picture of the rectangular specimens for the uniaxial tensile test; (c) a picture of the three different solutions for decellularization (Triton X-100, SDS, TnBP), plus the physiological solution.

### 2.2. Decellularization protocol and preparation of decellularized FL specimens

The chemical protocols for FL decellularization were designed based on the architecture of the native tissue, as suggested by Blaudez et al.^1^, to obtain suitable levels of DNA removal without causing significant damages to the ECM. The chemical agents employed in this study for the decellularization of FL tissues were Triton-X100, SDS, and TnBP. For each agent, we considered a total of six FL samples for the mechanical investigation (four were tested in February 2022 and two in May 2022, see Table S2 in the Supplementary Material). All samples were soaked in three different solutions (Figure 1c), in combination with 0.1% ethylenedinitrilotetraacetic acid (EDTA) and deionized water for 24 hours at room temperature (20 °C): (1) 1% solution of Triton X-100; (2) 0.1% solution of SDS; (3) 1% solution of TnBP. Afterwards, all specimens underwent 2 cycles of PBS (phosphate buffered saline) washes for 30 minutes. The creation of rectangular specimens for the mechanical tests (Figure 1b), as well as the measurement of their geometrical characteristics, were carried out following the same approach reported above.

### 2.3. Uniaxial tensile tests

All specimens were tested under uniaxial tensile tests to evaluate their macro-mechanical properties (Figure 2a). The tests were performed using an MTS Insight^®^ Electromechanical Testing Systems tensile machine, with a 1000 N load cell and sensitivity of 2.164 mV/V. All specimens were installed into 200N-loaded pneumatic clamps with knurling, and small gauze pads were inserted between the FL surfaces and the clamps in order to absorb the liquid released by the tissue, increase friction, and reduce the risk of slippage. The tensile tests were carried out using displacement control conditions, with a displacement rate of 0.1 mm/s and a sampling frequency of 30 Hz. The Young’s (elastic) modulus *E* was calculated as the slope of the linear elastic region in the stress-strain curves. The initial and final values of the linear elastic region were automatically detected by the TESTWORKS^®^ software of the testing machine. As mentioned above, the initial length of the specimens *L*_*0*_, used to compute the average engineering strain, was considered as the initial distance between the two clamps (see Figure 2b). After the tensile tests, macro-mechanical properties such as elastic modulus *E* and maximum stress *σ*_*max*_ were evaluated also considering propagation of uncertainties from the geometrical dimensions (see Supplementary Material).

**Figure 2.**
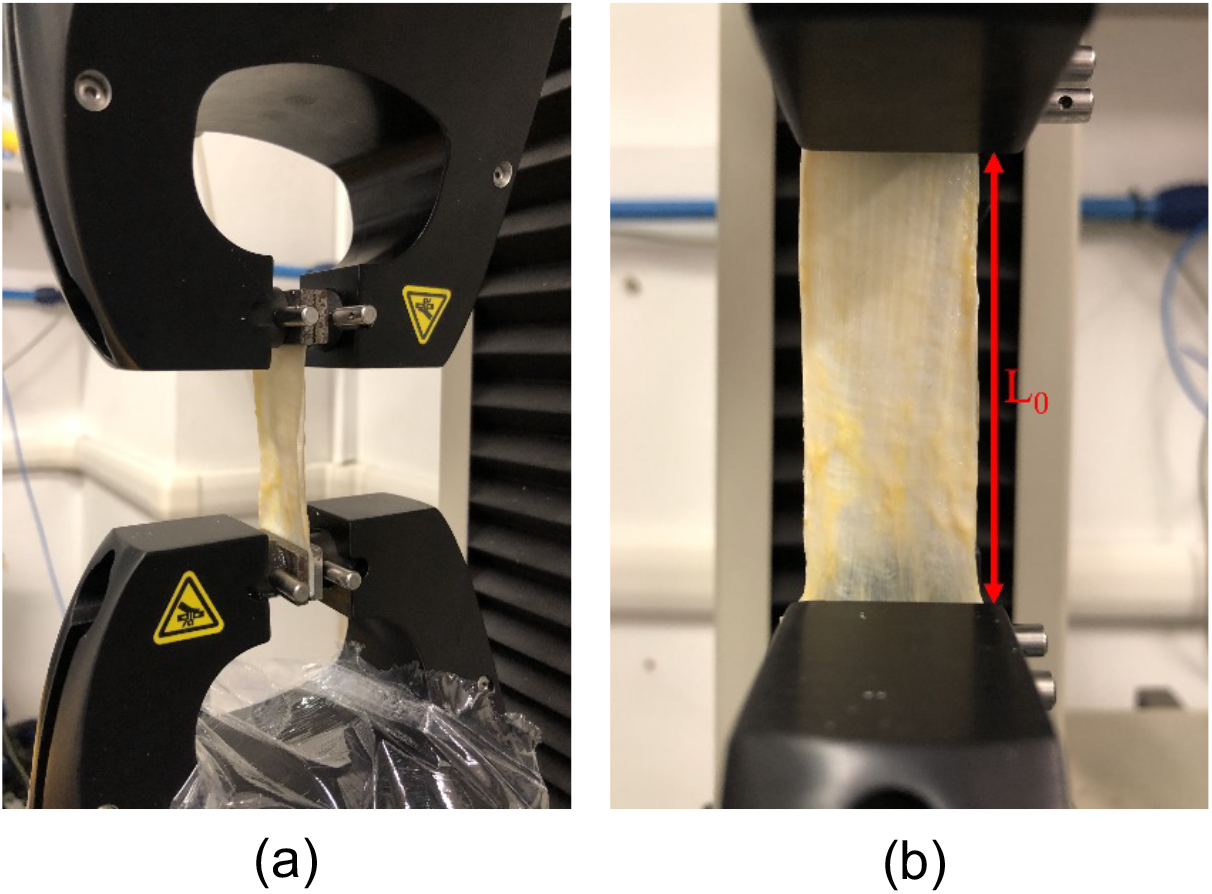
(a) Uniaxial tensile test of a FL sample; (b) measurement of the initial distance *L*_*0*_.

### 2.4. Digital Image Correlation (DIC) analysis

In combination with the tensile tests, Digital Image Correlation (DIC) was used to monitor the strain distribution in a few samples. For the DIC analysis, we investigated the displacement fields in Nat J4, TRX F4, SDS F3, and TnBP F3 specimens (see Tables S1 and S2 in the Supplementary Material for specimen labels). The DIC method was implemented with the VIC-EDU System, an optical non-contact method that can measure surface displacements. This enables the visualization of strain maps and deformation vector fields in all the directions, thus granting the characterization of inhomogeneous and anisotropic responses in complex specimens such as biological tissues^27–30^. A random paint pattern was applied on each sample (Figure 3a), by using a speckled pattern tool embedded with black water-based ink. The speckle size was chosen to fit in a desirable range of 1-5 pixels^27^. Subsequently, a series of images of the specimen undergoing uniaxial deformation was acquired. The software VIC-3D was used to calculate the strains between any two points within the area covered by the speckle pattern using virtual extensometers placed along the surface.

**Figure 3.**
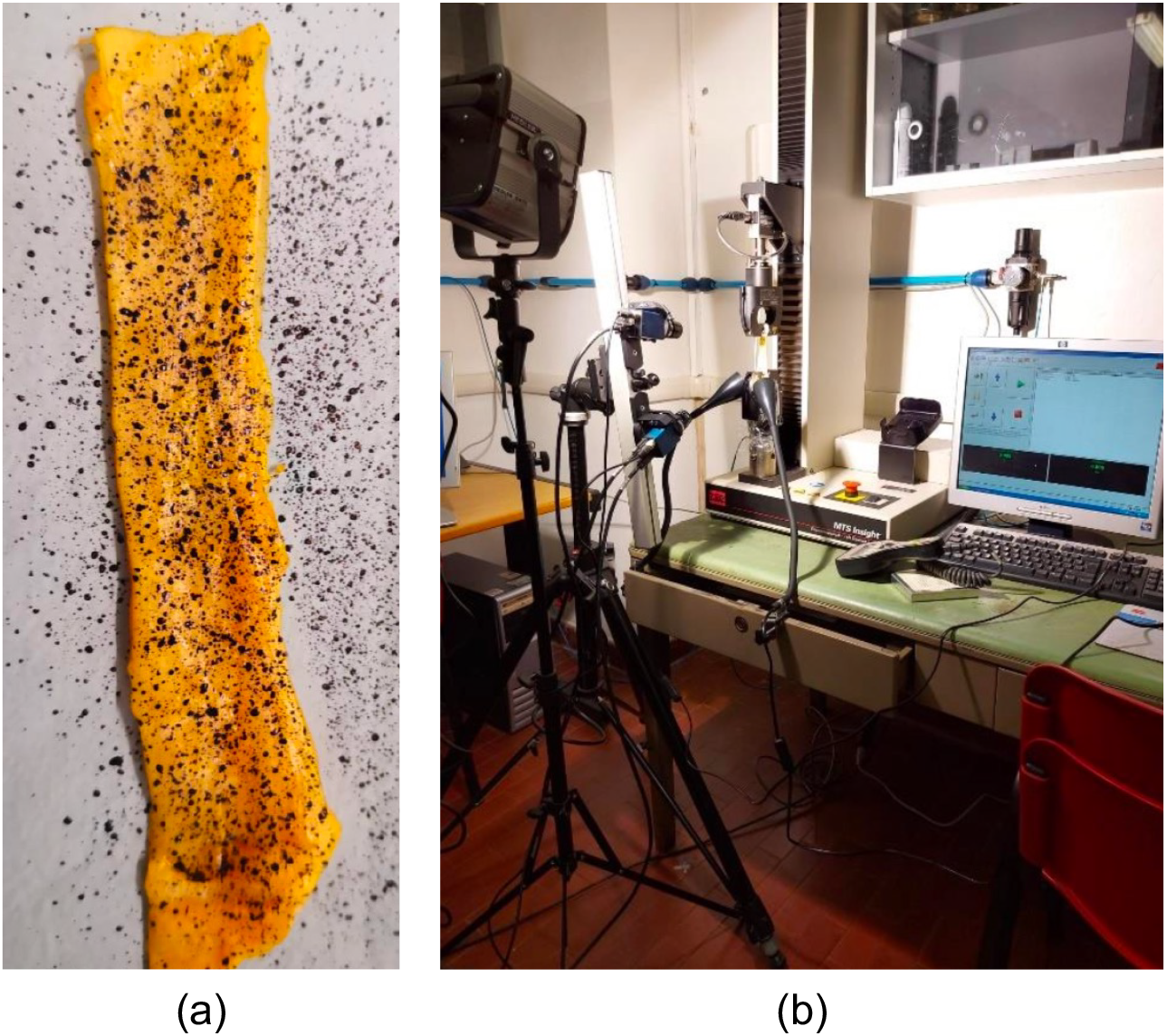
(a) Speckle pattern on a FL specimen; (b) integrated tensile test and DIC experimental setup.

The VIC-EDU System was placed 54.5 cm away from the specimen (Figure 3b). A three-dimensional configuration was used to compensate for possible out-of-plane displacements during testing. For better lighting of the specimen surface, two lights were added to the LED light integrated in the system. The VIC-Snap acquisition software was used to monitor the view field of the cameras and verify the good placement of the specimen and optimal lighting. The position of the lights was changed until the optimal lighting (50 ms of exposure time) was verified for all tested specimens. Images were acquired with two monochrome cameras, FLIR Blackfly^®^ U3-23S6M-C, with a progressive scan CMOS 1/1.2” image sensor, SONY IMX249. The resolution of the acquisition was 2.3 Megapixels, 1920 × 1200 pixels, with 5.86 × 5.86 μm pixel size. The system was calibrated by taking 30 images of the pattern calibration plate with different positions and orientations. During the test, both cameras were set to capture an image either every 2 or 3 seconds. For the post-processing in VIC-3D, an area of interest was defined for each sample and divided into smaller subsets of a size large enough to contain sufficient pattern information with minimal overlap. Due to the small dimension of the speckle, the size of the subsets was set to 9 and the step size that controls how many pixels are apart from each data point was set to 1. The reference point to enable the full-field displacement correlation was finally set in a position close to the lower grips.

## 3. Results and Discussion

### 3.1. Uniaxial tensile tests

The results of the mechanical tensile tests of all FL samples are reported in Tables S3-S5 in the Supplementary Material. Figure S1 shows the full force-displacement curves obtained by the testing machine for all specimens divided per treatment type, i.e., native, treated with Triton X-100, SDS, and TnBP. All samples were found to exhibit a pronounced softening behavior in the post-peak regime with no fragile collapse: after reaching the maximum load all samples allowed to accommodate higher displacements for lower load intensities before reaching ultimate failure (Figure S1). While for several specimens this was found to occur because of the capability of the tissue to redistribute tensile stresses across the collagen fiber network after the first fibers start to break, for some samples this was a consequence of large deformations/lacerations close to the machine clamps, highlighting a weakening of the tissue regions close to the external constrains. Force-displacements curves were then converted into stress-strain relationships, by considering the geometrical properties of each sample (cross-sectional area *A* and initial length *L*_*0*_), and are reported in Figure 4. Since a comparison in terms of stresses and strains in the post-peak regime is meaningless for our purposes, for clarity Figure 4 reports the stress-strain curves only up to the maximum stress *σ*_*max*_. A large variability across the FL samples was observed both in terms of stresses, strains, and stiffnesses (slopes of the curves), which is a typical fingerprint of the intrinsic variability of biomaterials^16,32^.

**Figure 4.**
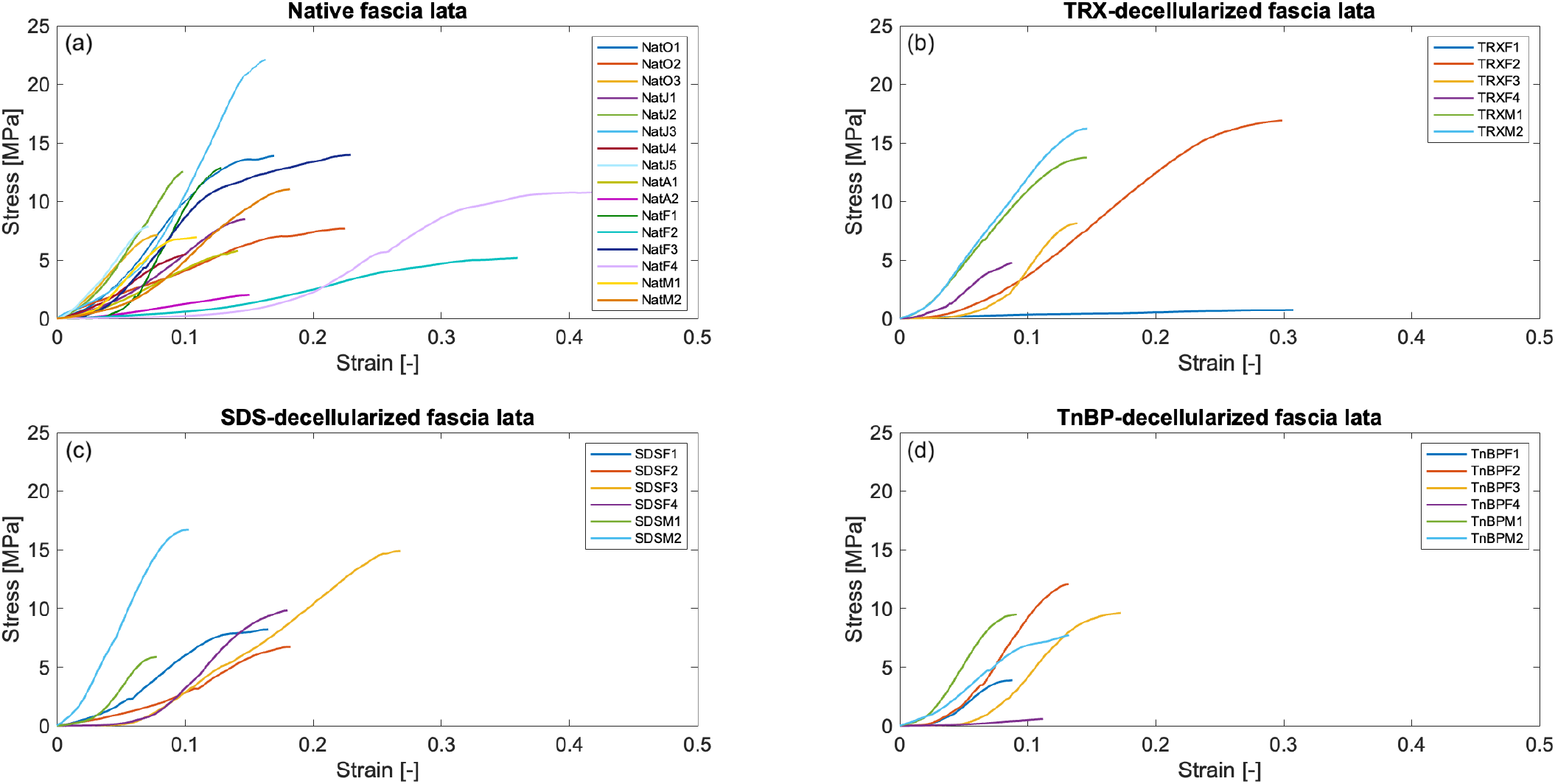
Stress-strain curves (up to *σ*_*max*_) for all FL specimens: (a) native samples; (b) samples treated with Triton X-100; (c) treated with SDS; (d) treated with TnBP. The colors of the curves refer to the different samples investigated during the various experimental campaigns (see Tables S1-S5 in the Supplementary Material for details about specimen labels). The full force-displacement curves, also including the post-peak regime, are reported in Figure S1.

Ensuring a proper mechanical rigidity for small deformations is pivotal to redistribute mechanical loads in physiological conditions, and the elastic modulus is a straightforward metric to quantify the material rigidity in the small-deformation regime. Figure 5a shows a comparison between the computed elastic moduli *E* across the population of all FL samples, colored according to the treatment protocol (native FL, treated with Triton X-100, SDS, TnBP). The elastic modulus was found to range from a minimum of ∼4 MPa to a maximum of ∼250 MPa, with an average value of ∼115 MPa. Despite the large variations, these values are in agreement with previous investigations on human samples^16,22,24,25^. A similar variability was also found for maximum stresses *σ*_*max*_, whose values are reported in Figure 5b. These were found to range from a minimum of ∼0.6 MPa to a maximum of ∼22 MPa, with an average value of ∼10 MPa. Also these values are in agreement with some previous analyses^16,25^. A clear positive correlation was observed between these two macro-mechanical properties, *E* and *σ*_*max*_ (see Figure S2 in the Supplementary Material), i.e., we generally found that stiffer samples (higher *E*) are also the ones that are able to withstand higher loads (higher *σ*_*max*_). A Pearson correlation coefficient between *E* and *σ*_*max*_ values of ∼0.7 was obtained across the whole dataset, with significant correlations also at the level of individual treatment groups, ranging from a minimum of ∼0.5 for SDS samples up to ∼0.9 for TnBP ones (see Figure S2 caption).

**Figure 5.**
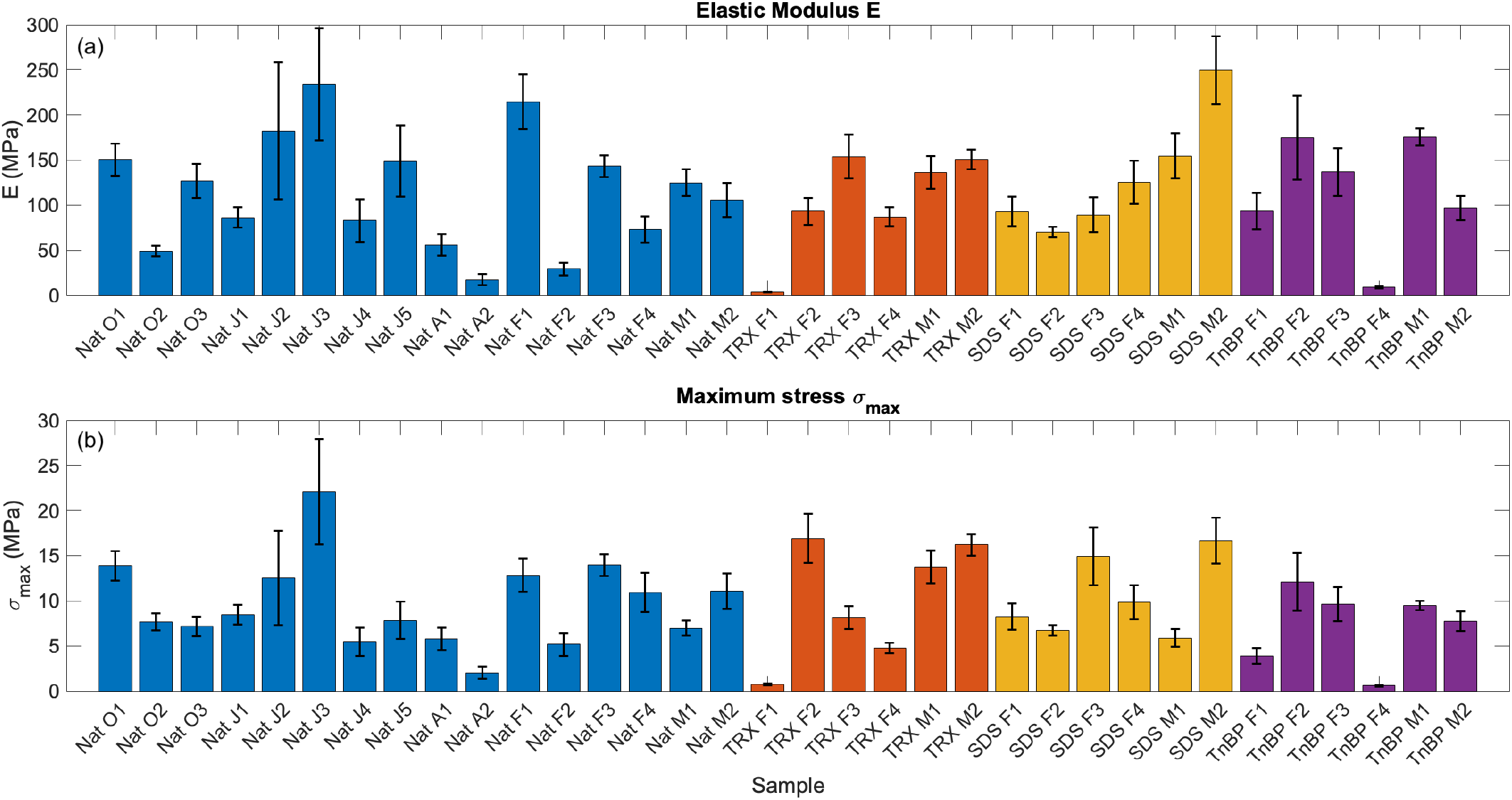
Comparison between (a) elastic moduli *E* and (b) maximum stresses *σ*_*max*_ for all FL samples (native FL group in blue, treated with Triton X-100 in red, treated with SDS in yellow, treated with TnBP in purple). For each sample, average and standard deviation values are shown (according to the values reported in Table S5 in the Supplementary Material).

To investigate the impact of the chemical treatment on the macro-mechanical properties of FL samples, we computed group-aggregated values for each FL group (native FL, treated with Triton X-100, SDS, TnBP). This is expected to average out the differences at the level of individual specimens yet taking into account the uncertainties that affects the measurement in each sample (see Supplementary Material). The results of these calculations considering the entire dataset of 34 samples seem to suggest a pronounced loss of mechanical performance when using Triton X-100 and TnBP treatments (see Table S7 and Figures S3a-S4a in the Supplementary Material). However, this is not directly observable from the data reported in Figure 5 (as well as in Table S5 and Figure S2 in the Supplementary Material), but most likely an effect coming from the low mechanical performance detected in only two samples, i.e., TRX F1 and TnBP F4, which with high probability are outliers of the *E* and *σ*_*max*_ distributions (see the extremely flat stress-strain curves of TRX F1 and TnBP F4 in Figure 4). To confirm the outlier nature of these specimens, as well as to remove the influence of all outliers on group-aggregated averages, we calculated 5^th^ and 95^th^ percentiles of the distributions of *E* and *σ*_*max*_ across all samples and considered outliers all specimens exceeding these boundaries. Table S6 in the Supplementary Material reports the list of lower- and upper-limit outliers, confirming that TRX F1 and TnBP F4 are lower-limit outliers for both the *E* and *σ*_*max*_ distribution. We also identified three additional upper-limit outliers (Table S6), i.e., Nat J3 (>95^th^ percentile of both *E* and *σ*_*max*_), SDS M2 (>95^th^ percentile of *E*), and TRX F2 (>95^th^ percentile of *σ*_*max*_). By removing these outliers from the group-aggregated calculations, elastic moduli and maximum stresses were found to exhibit more comparable values among the different treatment groups (see Figures S3-S4 in the Supplementary Material). Figure 6a reports aggregated values for the elastic moduli of each treatment group not considering the outliers (Nat J3, TRX F1, TRX F2, SDS M2, TnBP F4). In this case, we clearly observed that none of the decellularization method leads to a degradation of the tissue stiffness, with elastic moduli even above the average value in native samples (Figure 6a and Figures S3b-d). Similarly, Figure 6b reported the comparison in terms of *σ*_*max*_ values, showing that all treatment methods lead to a similar value of ultimate stress, around ∼7 MPa (see also Figure S4b-d). These results suggest that, on average, none of the decellularization protocols investigated here (Triton X-100, SDS, TnBP) leads to a mechanical degradation of the tissue, implying that these treatments can be effectively used - at least from a mechanical point of view - in surgical reconstruction applications.

**Figure 6.**
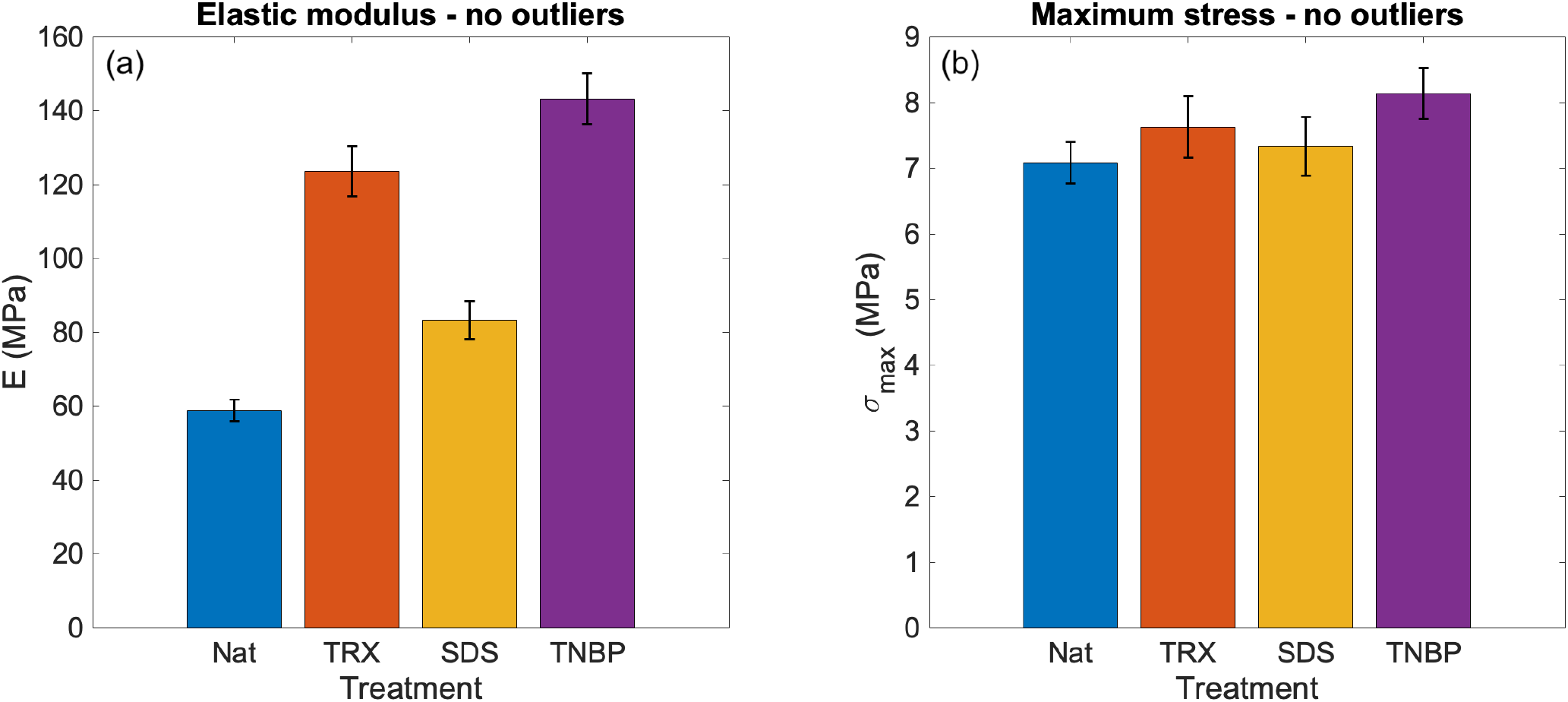
Values of (a) elastic moduli *E* and (b) ultimate stresses *σ*_*max*_, aggregated over each treatment group (native FL group in blue, Triton X-100 in red, SDS in yellow, TnBP in purple). Average values and error bars of elastic moduli have been computed according to Eqs. (S7a-b) in the Supplementary Material and considering all samples except for the outliers (5^th^ and 95^th^ percentiles) in both *E* and *σ*_*max*_ distributions (see Table S6 in the Supplementary Material). Figures S3-S4 in the Supplementary Material show treatment-aggregated quantities for both *E* and *σ*_*max*_ values with different outlier groups considered.

Nevertheless, a few limitations of the current experimental investigation need to be pointed out. First, the above conclusion holds true if outliers are removed from our group-aggregated calculations, especially the two samples TRX F1 and TnBP F4 which were found to exhibit remarkably low values of *E* and *σ*_*max*_. Note that TRX F2, TRX F3, and TRX F4 come from the same batch as TRX F1, and TnBP F1, TnBP F2, TnBP F3 from the same batch as TnBP F4. Therefore, the most likely explanation for such outlier behavior relies on the presence of microscopic defects in the ECM and fiber network of these specimens, which made the specific pieces of tissue weaker than the remaining portions. In *in vivo* conditions, localized weaknesses in specific areas of the tissue tend to be compensated by stronger and more rigid portions, redistributing stress concentrations over longer distances and reducing the risks of compliant mechanical responses. On the contrary, localized defects tend to emerge much more in laboratory setups, where the samples are only sub-mm thick and have a width of just a few dozen millimeters (Tables S1-S2). Secondly, our analysis would certainly benefit from a larger number of FL samples and a deeper statistical analysis. However, coming from human donors, the number of FL samples suitable for research purposes was extremely limited and highly unsteady over time. This left us with a relatively small population of samples. With a larger number of specimens, especially for each decellularization group, a more comprehensive statistical analysis could have been carried out, providing additional support for the identification of outliers and a more thorough comparison of treatment-aggregated properties. Lastly, material properties such as *E* and *σ*_*max*_ can help to describe the mechanical behavior at the macroscopic scale of the sample and, for this reason, are often extensively used in the literature^16,20–23,25^. Yet, both suffer from two main shortcomings: (1) they rely on over-simplifications of the irregular geometries and mechanical inhomogeneities, non-linearities, and anisotropies of the specimen, and (2) they do not allow to appreciate differences in the mechanical behavior at a smaller mesoscopic scale. With the employment of DIC, these limitations can be partially overcome, and the mechanical behavior of the FL tissue can be investigated at a smaller scale, allowing to highlight also localized effects up to ultimate collapse.

### 3.2. Digital Image Correlation (DIC)

To better assess the variations in strain patterns, strain deformations with DIC were quantified across a large rectangular area on the sample surface as well as within three smaller sub-regions (see Figure 7): an area close to the upper clamp (R_0_-Top), close to the specimen midpoint (R_1_-Center), and close to the lower clamp (R_2_-Down). Furthermore, two virtual strain gauges were employed to capture directional strain components: E_1_, aligned parallelly to the loading (longitudinal) direction and the collagen fibers, and E_0_, aligned in the perpendicular (transverse) direction.

**Figure 7.**
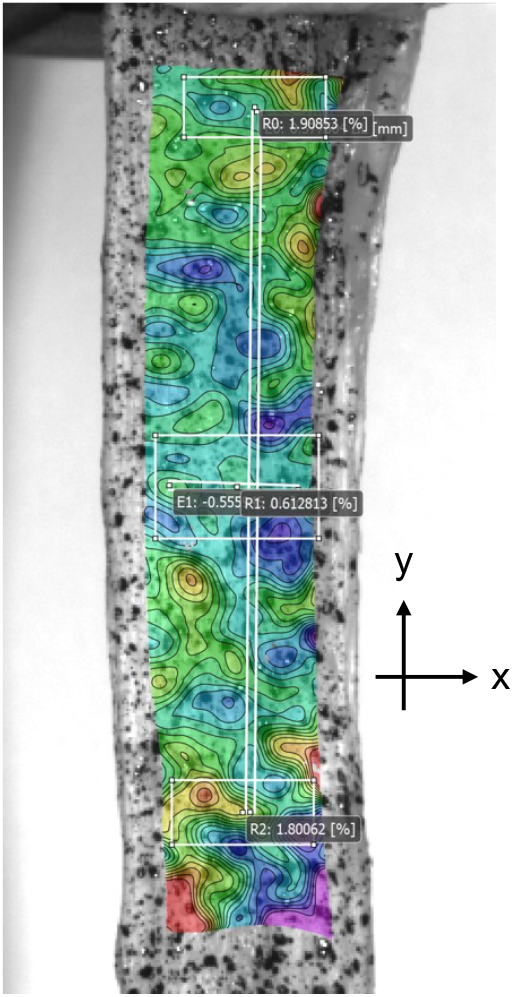
Example of a DIC strain map from one of the samples and position of the three smaller sub-regions (R_0_-Top, R_1_-Center, R_2_-Down). The colors of the DIC map reflect the intensities of the measured strains. The x-y reference system for the strain calculation is also reported. The two virtual strain gauges, E_1_ and E_0_, used to measure longitudinal and transversal deformations respectively, are also reported.

Figure 8 illustrates the evolution of DIC longitudinal strains (e_yy_) for the four investigated samples until maximum load *σ*_*max*_. For each sample, the four curves refer to the different focus regions (overall surface and upper, middle, and lower are). Table S8 in the Supplementary Material also reports maximum values of e_yy_, together with standard deviations evaluated across each sub-region. The native tissue (Nat J4) was found to exhibit an evolution of longitudinal DIC strains with maximum strain values ∼5% before collapse and higher strain localizations in the lower and upper regions (Figure 8a). The sample treated with Triton X-100 (TRX F4) showed a very similar deformative response compared to the native sample but with slightly lower strain values (∼4%) and a more homogeneous strain distribution across the three sub-regions (Figure 8b). Note that Nat J4 and TRX F4 were found exhibit very similar values of macro-mechanical properties (∼85 MPa for *E*, ∼5 MPa for *σ*_*max*_; see Table S5 in the Supplementary Material). The similar meso-mechanical response observed through DIC confirms the analogous mechanical-deformative behavior of these samples regardless of the chemical treatment. On the contrary, the sample treated with SDS (SDS F3) was found to exhibit a different behavior with an ultimate collapse occurring at larger DIC strains (∼15%) and a medium amount of inhomogeneity in the strain localizations (Figure 8c). Note that the elastic modulus of this sample was found to be very similar to the elastic moduli of Nat J4 and TRX F4 (∼85 MPa), while its ultimate stress was three times higher (∼15 MPa; see Table S5). This suggests that the SDS-treated sample has a similar elastic behavior as Nat J4 and TRX F4, but a higher capability to accommodate larger longitudinal strains before ultimate failure. Finally, the sample treated with TnBP (TnBP F3) was found to show an intermediate response compared to the three other specimens, with a strain evolution pattern similar to the native and TRX samples but at slightly higher strain values (∼7%, Figure 8d). Note that this sample was found to have a much higher elastic modulus (∼140 MPa), but a somewhat intermediate value of ultimate stress (∼10 MPa) compared to the other three.

**Figure 8.**
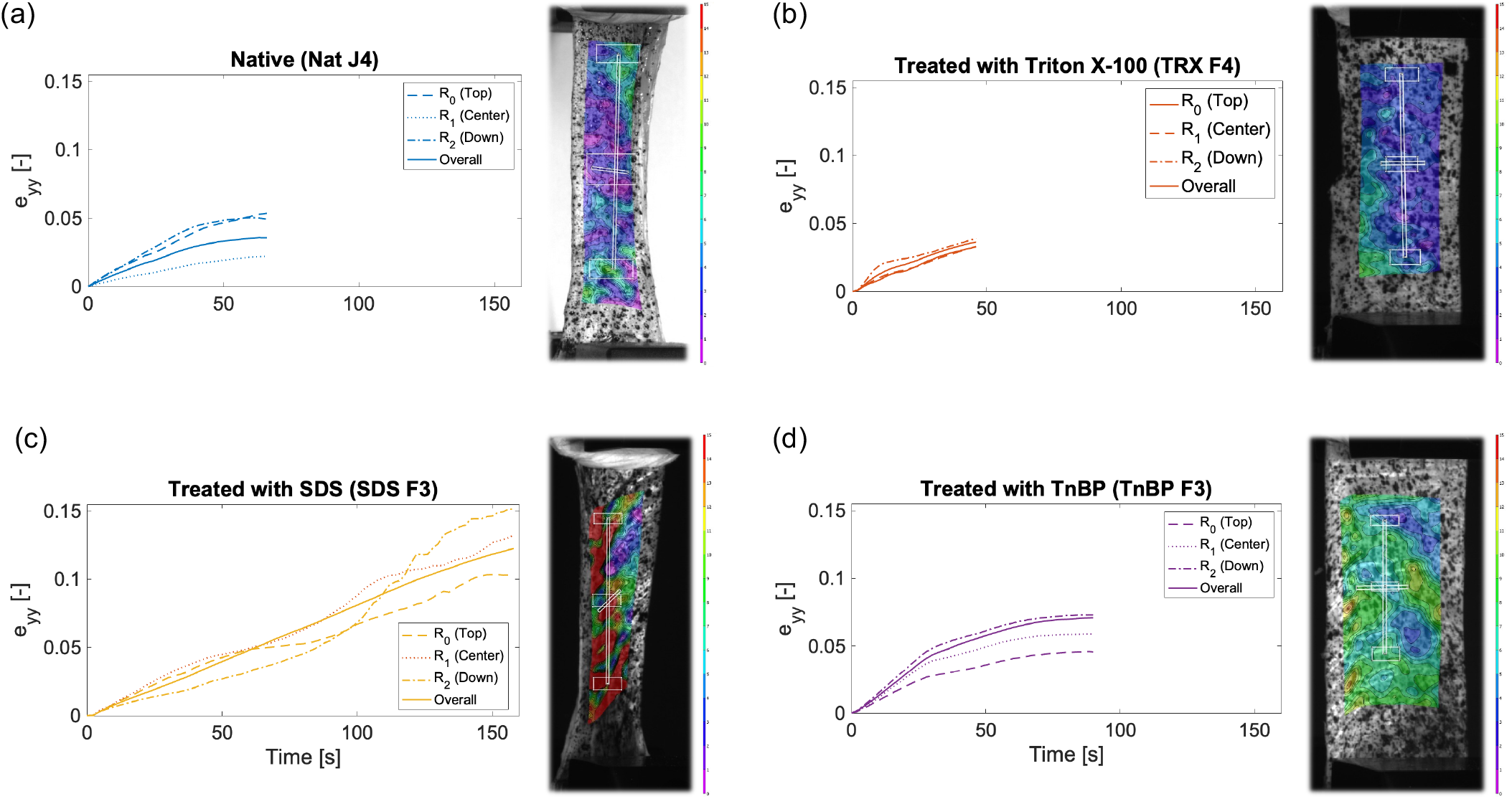
Time-evolution of DIC longitudinal strains (e_yy_) in: (a) native FL specimen (Nat J4), (b) sample treated with Triton X-100 (TRX F4), (c) sample treated with SDS (SDS F3), (d) sample treated with TnBP (TnBP F3). Each graph reports values of e_yy_ deformations evaluated on the overall surface (“Overall”) and on smaller sub-regions located in the upper (R_0_), middle (R_1_), and bottom (R_2_) portions of the specimen (see Figure 7). Snapshots of the DIC e_yy_ deformation fields are reported close to each graph.

Table S8 in the Supplementary Material summarizes not only maximum values of longitudinal strains (e_yy_), but also transversal (e_xx_) and Von Mises (e_VM_) deformations, with standard deviations across the averaging surfaces (overall, R_0_, R_1_, R_2_). These results highlight a few differences in the maximal mesoscopic deformations detected across the samples. The native FL sample shows Von Mises strains that are quite homogeneous across the specimen surface, with peaks at ∼11%. The sample treated with Triton X-100 displays maximum overall strains very similar to the native sample, with slightly lower spatial discrepancies (see also comparison between Figure 8a and Figure 8b), with Von Mises strains slightly lower compared to the native sample (∼7%). On the contrary, the sample treated with SDS undergoes larger maximum strains, both in the longitudinal and transversal direction (with peaks up to ∼18%) and Von Mises strains reaching values as high as ∼36%, confirming the larger deformations obtained in this sample. Finally, the sample treated with TnBP shows moderate strain values, with maximum Von Mises strains of ∼10%, confirming a somewhat intermediate deformation response of this sample (Table S8).

Based on the use of the two virtual gauges E_1_ and E_0_ (see Figure 7), we also evaluated the maximum longitudinal and transversal elongations of the specimen, as well as the Poisson ratios at the moment of maximum longitudinal elongation. As reported in Table S9 in the Supplementary Material, Poisson ratios were found to be ∼0.23 for the native specimen, ∼0.36 for TRX, ∼0.50 for SDS, and ∼0.20 for TnBP. While a few tests have been carried out to assess the anisotropic response of FL and highlighted the higher mechanical performance in the direction of collagen fibers^16,20,21,25^, to the best of our knowledge this is the first time the Poisson ratio of FL has been explicitly characterized - thanks to full-field DIC strain measurements. Given the highly inhomogeneous nature of FL specimens, we must emphasize that these values of Poisson ratios can only interpreted as preliminary estimates and would need further refinement with dedicated experimental tests. Yet, the fact that such different chemical treatments also lead to estimates of Poisson values of the same order of magnitude confirms that these decellularization protocols do not cause any significant alteration of the mechanical properties of FL.

Finally, it is interesting to compare longitudinal strains obtained from DIC to the average engineering strains evaluated from the stress-strain tensile curves at *σ*_*max*_ (see Figure 4). While the former were found to be ∼5% for NatJ4, ∼4% for TRX F4, ∼15% for SDS F3, ∼7% for TnBP F3, the latter were ∼10%, ∼8%, ∼25%, ∼18% (see Figure 4), respectively. These engineering values are in line with the strains at break reported in the literature^16^. Moreover, they hold the same relationship across different samples as obtained from DIC (TRX < Nat < TnBP < SDS), suggesting that we can use both DIC and standard tensile tests to evaluate the differences in the deformation properties of FL samples. Yet, in all samples the average engineering values tend to overestimate the strains obtained from full-field DIC measurements. This happens because the FL samples are not homogeneous, and we observed that strains can undergo intense redistributions within the sample volume, often localizing in the upper and lower extremities of the tissue in close proximity of the external constrains. We also observed the occurrence of localized deformations and lacerations close to the machine clamps, which explains why in the area monitored by DIC the strains exhibit lower values compared to the average engineering strains that are simply evaluated from the evolving distances between the machine clamps. Our results suggest that monitoring strains by full-field measurements can reveal strong inhomogeneities and localization effects, which cannot be appreciated by relying only on traditional average engineering strains. This in turn can affect the estimated values of macro-mechanical properties such as the elastic modulus. Hence, we suggest that further research efforts should be devoted at integrating the traditional stress measurements from the uniaxial tests to strain observations from DIC for a more reliable characterization of the tissue mechanical features.

## 4. Conclusions

In this work, we have carried out a mechanical characterization of FL samples from human dead donors to investigate their macro- and meso-mechanical properties under uniaxial tension. Both native samples and specimens treated with Triton X-100, SDS, and TnBP were investigated. From the uniaxial tests, we computed values of the ultimate strength *σ*_*max*_ and elastic modulus *E*. After removing outliers, we did not observe any significant deterioration of the (macro-mechanical) properties for any of the investigated decellularization protocols. We also used Digital Image Correlation (DIC) analysis to monitor the strain distribution within the specimen surfaces, both in the main direction of collagen fibers as well as in the orthogonal direction. The analysis of DIC strains indicated that the (meso-mechanical) deformation properties of FL specimens under different chemical treatments does not show extreme differences, leading to comparable strain evolutions and maximum strain values, as well as Poisson ratios of the same order of magnitude. The application of DIC analysis also proved effective in highlighting that relying on average engineering strains can lead to over- or under-estimations of the actual tissue deformations, especially when strain localizations, lacerations, or slippage events are found to occur close to the extremities^30^. Overall, the results of our analysis suggest that none of the investigated decellularization treatments deteriorates the mechanical performance of FL tissues, and they can therefore be used in the clinical practice, at least from a mechanical point of view. Further research efforts might be needed to increase the statistical significance of the mechanical characterization increasing the number of investigated samples, improve the experimental setup aiming to reduce the degree of lacerations close to the machine clumps, and integrate the DIC strain measurements with the stress measurements for a more accurate evaluation of stress-strain curves and macro-mechanical properties. Additional research on the histological and biophysical compatibility of the tissue upon decellularization would be also needed to validate the applicability of these chemical treatments in the clinical practice.

## Author Contributions

A.C. Faini, A. Leone, E. Camusso: Extraction of FL tissues from human dead donors; Preparation of the tissues; Application of the decellularization protocol. F. Demontis, G. Loi, E. Mazzotti, M. Reinoso Rodriguez, D. Scaramozzino: Preparation of FL tissues for the mechanical tests; Measuring of geometrical properties; Uniaxial tensile tests. F. Vecchio: DIC acquisition and post-processing. F. Genzano Besso, C. Surace, G. Lacidogna: Acquisition of funding and supervision. D. Scaramozzino: Project supervision; Post processing of results; Writing of the paper draft. All authors contributed to discussions and were involved in the draft revision.

## Data Availability

All the macro-mechanical data as well as movies of DIC strain maps are freely available on figshare at https://doi.org/10.6084/m9.figshare.28768406, https://doi.org/10.6084/m9.figshare.28768421, and https://doi.org/10.6084/m9.figshare.28845464.

## Acknowledgments

We acknowledge “Banca dei Tessuti Muscoloscheletrici di Torino” for providing the FL samples (resulted not eligible for transplant) from human donors, and the Department of Structural, Building, and Geotechnical Engineering (Politecnico di Torino) for providing the facility and equipment to carry out mechanical tests.

