## Supplementary Material for "Uniaxial tensile tests and Digital Image Correlation analysis for the mechanical characterization of human Fascia Lata under different decellularization treatments"

^1^Department of Structural, Geotechnical and Building Engineering, Politecnico di Torino, Torino, Italy. ^2^BIO MAterials & STructures (BIOMAST) Laboratory, Politecnico di Torino, Torino, Italy. ^3^Department of Medical Sciences, Università degli Studi di Torino, Torino, Italy. ^4^Banche dei Tessuti e Bioconservatorio, Azienda Ospedaliera Universitaria Città della Salute e della Scienza di Torino, Torino, Italy. ^5^Centro Regionale Trapianti, Azienda Ospedaliera Universitaria Città della Salute e della Scienza di Torino, Torino, Italy. ^6^Department of Oncology-Pathology, Karolinska Institutet, Solna, Sweden.

^$^These authors contributed equally.

**Supplementary Material**

Geometrical properties of the samples, error propagation and calculation of mechanical properties

As explained in the main text, the geometrical characteristics of the samples were measured via digital a digital caliber and a digital micrometer. The resting length of the sample, *L_0_*, was considered equal to the initial gripping distance in the testing machine at the beginning of the test (see Figure 2b in the main text) and used to compute average engineering strains. Since *L_0_* can be defined with high confidence, i.e., with an error of the order of ~µm due to the sensitivity of the digital micrometer, we did not consider any uncertainty for *L_0_* (Tables S1-S2). A larger variability was observed for the width *W* and thickness *T* of the FL samples, as these were not characterized by perfectly homogeneous geometries (see Figure 1b in the main text). We measured five distinct values of *W* and *T* along the sample surface and then computed average values *W_µ_* and *T_µ_* and standard deviations *W_σ_* and *T_σ_* as:

$$W_{\mu}=\frac{1}{5}\sum_{i=1}^{5} W_{i} W_{\sigma}=\sqrt{\frac{1}{4}\sum_{i=1}^{5} \left( W_{i}-W_{\mu} \right)^{2}}$$

(S1a)

$$T_{\mu}=\frac{1}{5}\sum_{i=1}^{5} T_{i} T_{\sigma}=\sqrt{\frac{1}{4}\sum_{i=1}^{5} \left( T_{i}-T_{\mu} \right)^{2}}$$

(S1b)

All width and thickness values for both native Fascia Lata (FL) samples are reported in Table S1 and S2, respectively. Average values and standard deviations were subsequently used to compute averages of functional variables that depend on these geometrical quantities as well as to propagate errors. This was done based on the classical error-propagation equations:

$$f=f(x_{1},x_{2},\ldots,x_{n}) \to f_{\mu}=f({x_{1}}_{\mu},{x_{2}}_{\mu},\ldots,{x_{n}}_{\mu}) f_{\sigma}=\sqrt{\sum_{i=1}^{n} \left( \frac{\partial f}{\partial x_{i}} \right)^{2}{{x_{i}}_{\sigma}}^{2}}$$

(S2)

where *f* is a generic function of the *n* variables *x_i_*, *f_µ_* represents its mean value, and *f_σ_* the related standard deviation error. Based on average and standard deviation values of *W* and *T*, the cross-sectional areas of each sample were estimated (Tables S1-S2). The average value of the cross-sectional area is simply given by the product between average values of width and thickness, while its standard deviation was computed by accounting for the propagation of width and thickness uncertainties, as:

$$A=W\times T \to A_{\mu}=W_{\mu}\times T_{\mu} A_{\sigma}=\sqrt{{T_{\mu}}^{2}{W_{\sigma}}^{2}+{W_{\mu}}^{2}{T_{\sigma}}^{2}}$$

(S3)

Based on the knowledge of geometrical dimensions and force/displacement outputs from the tensile machine, macro-mechanical properties could be computed for each sample, such as the maximum stress *σ_max_* and the elastic modulus *E*. The ultimate stress can be simply obtained as the ratio between the maximum force detected during the tensile test, *F_max_*, and the cross-sectional area *A*. For *F_max_*, we did not consider any measurement uncertainty, as the tensile machine has a sensitivity of ~mN, which is well below the intrinsic variability of force measurements in biological samples. On the other hand, the larger uncertainties in the values of cross-sectional area *A* were considered to propagate errors, as:

$$\sigma_{max}=\frac{F_{max}}{A} \to\sigma_{\mu}=\frac{F_{max}}{A_{\mu}} \sigma_{\sigma}=\frac{F_{max}}{{A_{\mu}}^{2}}A_{\sigma}$$

(S4)

The elastic modulus *E* was computed based on the knowledge of the limit points of the linear-elastic regions, *B* and *M*, automatically identified by the testing machine. Based on the force (*F_B_* and *F_M_*) and displacement values (*L_B_* and *L_M_*) at these points, *E* was simply computed as:

$$E=\frac{\Delta\sigma_{B-M}}{\Delta\varepsilon_{B-M}}=\frac{\Delta F_{B-M}/A}{\Delta L_{B-M}/L_{0}}= \frac{L_{0}\left( F_{M}-F_{B} \right)}{A\left( L_{M}-L_{B} \right)} \to E_{\mu}=\frac{L_{0}\left( F_{M}-F_{B} \right)}{A_{\mu}\left( L_{M}-L_{B} \right)} E_{\sigma}=\frac{L_{0}\left( F_{M}-F_{B} \right)}{{A_{\mu}}^{2}\left( L_{M}-L_{B} \right)}A_{\sigma}$$

(S5)

where, as usual, we considered the error propagation based on the main source of uncertainty, i.e., the geometrical dimension of the sample. Values of maximum forces and forces/displacements at the borders of the linear-elastic region for native and treated FL samples are reported in Tables S3 and S4, respectively. As a consequence, elastic moduli and maximum stresses, with their corresponding uncertainties, are reported in Table S5.

Calculation of group-aggregated mechanical properties and identification of outliers

The values of both elastic moduli and maximum stresses were found to exhibit a large variability, not only across different treatment groups, but also within the same group. For example, among the 16 FL native specimens, maximum stresses were found to range from a minimum of ~1 MPa to a maximum of ~22 MPa, and elastic moduli of the same group from a minimum of ~4 MPa to an astonishing maximum of ~235 MPa. Each of these values also carries different amount of measurement uncertainties due to the different standard deviation associated with geometrical properties of the samples (Tables S1-S5). To aggregate data consistently for each FL group (native, Triton X-100, SDS, TnBP) and consider all standard deviation values, we weighted each measurement *j* by the inverse of its corresponding variance, i.e.:

$$w_{\sigma_{max,j}}=\frac{1}{{\sigma_{{max,j}_{\sigma}}}^{2}} w_{E_{j}}=\frac{1}{{E_{j_{\sigma}}}^{2}}$$

(S6)

and then these weights were used to compute group-aggregated averages and standard deviations of maximum stresses and elastic moduli for each group of samples *G* as.:

$${\sigma_{max, G}}_{\mu}={\sum_{j\in G} {\sigma_{max, j}}_{\mu}w_{\sigma_{max,j}}}/{\sum_{j\in G} w_{\sigma_{max,j}}} {\sigma_{max, G}}_{\sigma}=\sqrt{1/{\sum_{j\in G} w_{\sigma_{max,j}}}}$$

(S7a)

$${E_{G}}_{\mu}={\sum_{j\in G} {E_{j}}_{\mu}w_{E_{j}}}/{\sum_{j\in G} w_{E_{j}}} {E_{G}}_{\sigma}=\sqrt{1/{\sum_{j\in G} w_{E_{j}}}}$$

(S7b)

where *G* is associated with the treatment group, i.e., “Nat”, “TRX”, “SDS”, or “TNBP”. In this way, measurements with the lowest variance end up having a larger weight in the definition of the group-aggregated values of maximum stresses and elastic moduli.

Calculations based on Eqs. (S6-7) have been carried out both on the whole dataset (see Figures S3a-S4a) and also considering subsets of samples not including outliers. As a matter of fact, from the analysis of elastic moduli and maximum stresses (Figure S2) as well as stress-strain curves (Figure 4 in the main text), it is evident how certain samples were found to exhibit a remarkably different mechanical behavior compared to the remaining population, e.g., TRX F1. To identify such outliers in a quantitative way, we looked at the whole population of elastic moduli *E* and maximum stresses *σ_max_*, and defined outliers those with values exceeding the 5^th^ and 95^th^ percentiles (Table S6). For the purpose of finding group-aggregated mechanical properties, these outliers were then removed from the dataset, and average values and standard deviations of elastic moduli and maximum stresses for each group were calculated again according to Eqs. (S6-7). Table S7 and Figures S3-S4 report group-aggregated values based on the selected dataset, i.e., all 34 samples, all samples except outliers of the *E* population, all samples except outliers of the *σ_max_* population, and all samples except outliers of both the *σ_max_* and *E* populations.

**Table S1.** Dimensions of native FL specimens. L_0_ stands for the initial length of the sample within the tensile test machine (see Figure 2b). The width W and thickness T of the samples are reported through average values and standard deviations. A denotes the cross-sectional area of the sample used for the calculation of the stresses. The labelling of native specimens (“Nat”) refers to the month of the experimental campaign (“O”: October 2020, “J”: January 2021, “A”: April 2021, “F”: February 2022, “M”: May 2022), and to the progressive identification number of each sample within each campaign.

| Specimen | L_0_ [mm] | W [mm] | T [mm] | A [mm^2^] |
| --- | --- | --- | --- | --- |
| Nat O1 | 33.24 | 14.17 ± 1.55 | 0.36 ± 0.02 | 5.07 ± 0.60 |
| Nat O2 | 28.83 | 14.66 ± 1.79 | 0.40 ± 0.01 | 5.86 ± 0.72 |
| Nat O3 | 44.60 | 15.43 ± 2.30 | 0.49 ± 0.01 | 7.61 ± 1.14 |
| Nat J1 | 69.66 | 17.24 ± 1.38 | 1.65 ± 0.16 | 28.41 ± 3.64 |
| Nat J2 | 80.35 | 15.14 ± 1.99 | 1.08 ± 0.43 | 16.38 ± 6.83 |
| Nat J3 | 68.15 | 14.74 ± 0.75 | 0.64 ± 0.17 | 9.39 ± 2.49 |
| Nat J4 | 64.98 | 18.24 ± 1.63 | 1.56 ± 0.42 | 28.50 ± 8.11 |
| Nat J5 | 65.28 | 20.05 ± 1.04 | 1.01 ± 0.26 | 20.23 ± 5.36 |
| Nat A1 | 60.73 | 12.66 ± 1.82 | 0.28 ± 0.04 | 3.54 ± 0.77 |
| Nat A2 | 72.47 | 16.71 ± 5.06 | 0.85 ± 0.14 | 14.17 ± 4.87 |
| Nat F1 | 51.93 | 24.90 ± 2.00 | 0.61 ± 0.07 | 15.27 ± 2.17 |
| Nat F2 | 22.69 | 22.13 ± 4.85 | 1.35 ± 0.14 | 29.81 ± 7.25 |
| Nat F3 | 39.56 | 22.85 ± 1.90 | 0.62 ± 0.01 | 14.19 ± 1.21 |
| Nat F4 | 18.34 | 23.78 ± 0.87 | 0.79 ± 0.15 | 18.74 ± 3.70 |
| Nat M1 | 50.95 | 17.17 ± 0.89 | 0.62 ± 0.07 | 10.68 ± 1.26 |
| Nat M2 | 38.77 | 16.96 ± 2.49 | 0.37 ± 0.04 | 6.28 ± 1.12 |

**Table S2.** Dimensions of decellularized FL specimens. L_0_, W, T, and A have the same meaning as in Table 1. The labelling of specimens reflects the chemical agent for the decellularization protocol (“TRX”: Triton X-100, “SDS”: SDS, “TNBP”: TnBP), the month of the experimental campaign (“F”: February 2022, “M”: May 2022), and the progressive identification number of each individual sample within each campaign.

| Specimen | L_0_ [mm] | W [mm] | T [mm] | A [mm^2^] |
| --- | --- | --- | --- | --- |
| TrX F1 | 27.42 | 26.93 ± 1.16 | 0.31 ± 0.04 | 8.27 ± 1.03 |
| TrX F2 | 22.48 | 26.81 ± 1.34 | 0.59 ± 0.09 | 15.79 ± 2.51 |
| TrX F3 | 31.09 | 23.20 ± 1.32 | 0.60 ± 0.09 | 13.87 ± 2.19 |
| TrX F4 | 51.88 | 23.56 ± 0.78 | 0.35 ± 0.04 | 8.20 ± 1.00 |
| TrX M1 | 44.67 | 22.91 ± 1.43 | 0.83 ± 0.10 | 19.13 ± 2.52 |
| TrX M2 | 53.15 | 18.58 ± 0.82 | 0.47 ± 0.03 | 8.73 ± 0.63 |
| SDS F1 | 29.88 | 24.89 ± 1.71 | 0.58 ± 0.09 | 14.41 ± 2.54 |
| SDS F2 | 34.08 | 23.15 ± 0.22 | 0.70 ± 0.06 | 16.11 ± 1.40 |
| SDS F3 | 58.50 | 22.49 ± 2.04 | 0.51 ± 0.10 | 11.38 ± 2.45 |
| SDS F4 | 44.62 | 23.61 ± 3.09 | 0.51 ± 0.07 | 12.06 ± 2.29 |
| SDS M1 | 51.33 | 19.22 ± 0.90 | 0.66 ± 0.10 | 12.70 ± 2.07 |
| SDS M2 | 52.67 | 19.27 ± 2.14 | 0.84 ± 0.09 | 16.11 ± 2.45 |
| TnBP F1 | 34.74 | 25.88 ± 0.37 | 0.83 ± 0.18 | 21.38 ± 4.72 |
| TnBP F2 | 39.35 | 22.87 ± 0.50 | 0.81 ± 0.21 | 18.43 ± 4.89 |
| TnBP F3 | 51.45 | 24.21 ± 1.12 | 0.45 ± 0.09 | 10.99 ± 2.14 |
| TnBP F4 | 49.17 | 16.31 ± 2.37 | 0.35 ± 0.04 | 5.76 ± 1.09 |
| TnBP M1 | 50.48 | 21.47 ± 0.73 | 0.56 ± 0.02 | 12.00 ± 0.64 |
| TnBP M2 | 45.51 | 21.58 ± 1.35 | 0.30 ± 0.04 | 6.45 ± 0.89 |

**Table S3.** Maximum force F_max_ and values of force (F_B_ and F_M_) and displacements (L_B_ and L_M_) at the limits of the linear-elastic region for the 16 native FL specimens. We also report the stiffness S (in N/mm), which is computed as the ratio between the increment of force ΔF vs. increment of displacement ΔL within the linear elastic region.

| Specimen | F_max_ [N] | F_B_ [N] | F_M_ [N] | L_B_ [mm] | L_M_ [mm] | ΔF [N] | ΔL [mm] | S [N/mm] |
| --- | --- | --- | --- | --- | --- | --- | --- | --- |
| Nat O1 | 70.46 | 27.06 | 41.43 | 2.21 | 2.84 | 14.37 | 0.63 | 22.96 |
| Nat O2 | 45.11 | 25.67 | 34.94 | 3.07 | 4.00 | 9.28 | 0.93 | 9.93 |
| Nat O3 | 54.34 | 16.74 | 27.94 | 1.25 | 1.77 | 11.20 | 0.52 | 21.57 |
| Nat J1 | 240.95 | 132.32 | 181.11 | 6.29 | 7.68 | 48.80 | 1.39 | 35.21 |
| Nat J2 | 205.35 | 129.63 | 171.24 | 5.49 | 6.61 | 41.61 | 1.12 | 37.15 |
| Nat J3 | 207.33 | 134.61 | 176.73 | 7.97 | 9.28 | 42.12 | 1.31 | 32.25 |
| Nat J4 | 155.88 | 63.48 | 95.51 | 3.15 | 4.03 | 32.02 | 0.88 | 36.43 |
| Nat J5 | 158.63 | 89.15 | 121.17 | 2.76 | 3.45 | 32.02 | 0.69 | 46.13 |
| Nat A1 | 20.53 | 7.41 | 11.59 | 2.28 | 3.57 | 4.18 | 1.28 | 3.26 |
| Nat A2 | 28.75 | 7.54 | 13.69 | 2.19 | 4.00 | 6.15 | 1.81 | 3.39 |
| Nat F1 | 196.16 | 40.52 | 80.04 | 0.63 | 1.26 | 39.52 | 0.63 | 63.13 |
| Nat F2 | 154.03 | 74.41 | 106.01 | 1.90 | 2.73 | 31.60 | 0.83 | 38.21 |
| Nat F3 | 198.21 | 82.41 | 122.19 | 1.55 | 2.32 | 39.77 | 0.77 | 51.38 |
| Nat F4 | 205.11 | 107.41 | 147.13 | 1.48 | 2.01 | 39.73 | 0.53 | 74.53 |
| Nat M1 | 74.30 | 27.77 | 43.14 | 1.06 | 1.64 | 15.37 | 0.59 | 26.18 |
| Nat M2 | 69.39 | 27.62 | 41.77 | 1.61 | 2.43 | 14.16 | 0.83 | 17.14 |

**Table S4.** Maximum force F_max_ and values of force (F_B_ and F_M_) and displacements (L_B_ and L_M_) at the limits of the linear-elastic region for the 18 FL specimens that underwent decellularization treatments. We also report the stiffness S (in N/mm), which is computed as the ratio between the increment of force ΔF vs. increment of displacement ΔL within the linear elastic region.

| Specimen | F_max_ [N] | F_B_ [N] | F_M_ [N] | L_B_ [mm] | L_M_ [mm] | ΔF [N] | ΔL [mm] | S [n/mm] |
| --- | --- | --- | --- | --- | --- | --- | --- | --- |
| TrX F1 | 6.08 | 0.49 | 1.73 | 0.40 | 1.49 | 1.24 | 1.09 | 1.13 |
| TrX F2 | 267.23 | 117.04 | 171.08 | 1.81 | 2.64 | 54.04 | 0.83 | 65.34 |
| TrX F3 | 112.88 | 45.75 | 69.59 | 0.66 | 1.01 | 23.84 | 0.35 | 68.69 |
| TrX F4 | 39.04 | 10.15 | 18.60 | 0.59 | 1.21 | 8.45 | 0.61 | 13.76 |
| TrX M1 | 262.97 | 90.61 | 120.20 | 1.55 | 2.06 | 29.59 | 0.51 | 58.36 |
| TrX M2 | 141.55 | 20.66 | 50.42 | 0.82 | 2.02 | 29.76 | 1.20 | 24.80 |
| SDS F1 | 118.67 | 33.67 | 57.55 | 0.76 | 1.29 | 23.88 | 0.53 | 44.80 |
| SDS F2 | 108.47 | 50.95 | 73.03 | 1.44 | 2.11 | 22.08 | 0.67 | 33.15 |
| SDS F3 | 169.71 | 112.99 | 146.28 | 6.56 | 8.48 | 33.29 | 1.92 | 17.34 |
| SDS F4 | 118.71 | 50.12 | 74.54 | 1.43 | 2.15 | 24.42 | 0.72 | 33.92 |
| SDS M1 | 74.71 | 36.24 | 51.55 | 0.94 | 1.34 | 15.31 | 0.40 | 38.28 |
| SDS M2 | 269.10 | 119.62 | 174.58 | 1.55 | 2.27 | 54.95 | 0.72 | 76.32 |
| TnBP F1 | 83.24 | 36.98 | 54.55 | 0.64 | 0.95 | 17.57 | 0.31 | 57.42 |
| TnBP F2 | 222.44 | 76.18 | 121.99 | 0.92 | 1.48 | 45.81 | 0.56 | 81.81 |
| TnBP F3 | 105.89 | 40.96 | 62.78 | 1.37 | 2.12 | 21.82 | 0.75 | 29.21 |
| TnBP F4 | 3.45 | 1.28 | 2.16 | 1.22 | 2.04 | 0.88 | 0.83 | 1.06 |
| TnBP M1 | 113.96 | 54.02 | 77.39 | 1.27 | 1.83 | 23.37 | 0.56 | 41.73 |
| TnBP M2 | 49.88 | 19.23 | 30.96 | 1.39 | 2.25 | 11.72 | 0.85 | 13.73 |

**Table S5.** Mechanical properties (Young’s moduli and maximum stresses) for the 16 native FL samples and the 18 decellularized specimens. The elastic moduli and maximum stresses are reported with their average and standard deviation values that account for the propagation of uncertainties in geometrical quantities.

| Specimen | E [MPa] | σ_max_ [MPa] | Specimen | E [MPa] | σ_max_ [MPa] |
| --- | --- | --- | --- | --- | --- |
| Nat O1 | 150.41 ± 17.81 | 13.89 ± 1.64 | TrX F1 | 3.76 ± 0.47 | 0.74 ± 0.09 |
| Nat O2 | 48.83 ± 5.97 | 7.69 ± 0.94 | TrX F2 | 93.02 ± 14.81 | 16.92 ± 2.69 |
| Nat O3 | 126.49 ± 18.89 | 7.14 ± 1.07 | TrX F3 | 153.96 ± 24.30 | 8.14 ± 1.28 |
| Nat J1 | 86.33 ± 11.06 | 8.48 ± 1.09 | TrX F4 | 87.05 ± 10.65 | 4.76 ± 0.58 |
| Nat J2 | 182.18 ± 75.96 | 12.53 ± 5.22 | TrX M1 | 136.25 ± 17.98 | 13.74 ± 1.81 |
| Nat J3 | 234.13 ± 62.16 | 22.09 ± 5.86 | TrX M2 | 150.92 ± 10.91 | 16.20 ± 1.17 |
| Nat J4 | 83.07 ± 23.64 | 5.47 ± 1.56 | SDS F1 | 92.89 ± 16.38 | 8.23 ± 1.45 |
| Nat J5 | 148.86 ± 39.42 | 7.84 ± 2.08 | SDS F2 | 70.11 ± 6.08 | 6.73 ± 0.58 |
| Nat A1 | 55.91 ± 12.07 | 5.79 ± 1.25 | SDS F3 | 89.16 ± 19.23 | 14.91 ± 3.22 |
| Nat A2 | 17.33 ± 5.96 | 2.03 ± 0.70 | SDS F4 | 125.44 ± 23.76 | 9.84 ± 1.86 |
| Nat F1 | 214.74 ± 30.55 | 12.85 ± 1.83 | SDS M1 | 154.68 ± 25.17 | 5.88 ± 0.96 |
| Nat F2 | 29.09 ± 7.07 | 5.17 ± 1.26 | SDS M2 | 249.56 ± 37.96 | 16.71 ± 2.54 |
| Nat F3 | 143.24 ± 12.18 | 14.00 ± 1.19 | TnBP F1 | 93.30 ± 20.60 | 3.89 ± 0.86 |
| Nat F4 | 72.95 ± 14.41 | 10.95 ± 2.16 | TnBP F2 | 174.64 ± 46.31 | 12.07 ± 3.20 |
| Nat M1 | 124.93 ± 14.77 | 6.96 ± 0.82 | TnBP F3 | 136.71 ± 26.66 | 9.63 ± 1.88 |
| Nat M2 | 105.86 ± 18.83 | 11.06 ± 1.97 | TnBP F4 | 9.08 ± 1.72 | 0.60 ± 0.11 |
|  |  |  | TnBP M1 | 175.55 ± 9.36 | 9.50 ± 0.51 |
|  |  |  | TnBP M2 | 96.82 ± 13.42 | 7.73 ± 1.07 |

**Table S6.** Outliers exceeding the 5^th^ and 95^th^ percentiles of the E distribution, the σ_max_ distribution, or both.

|  | *E* distribution | *σ_max_* distribution | *σ_max_* + *E* distributions |
| --- | --- | --- | --- |
| Lower-limit outliers (<5^th^ percentile) | TRX F1, TnBP F4 | TRX F1, TnBP F4 | TRX F1, TnBP F4 |
| Upper-limit outliers (>95^th^ percentile) | Nat J3, SDS M2 | Nat J3, TRX F2 | Nat J3, TRX F2, SDS M2 |
| All outliers | Nat J3, TRX F1, SDS M2, TnBP F4 | Nat J3, TRX F1, TRX F2, TnBP F4 | Nat J3, TRX F1, TRX F2, SDS M2, TnBP F4 |

**Table S7.** Values of elastic modulus E and maximum stress σ_max_ for each FL group, i.e., native, treated with Triton X-100, SDS, or TnBP, considering all investigated specimens or excluding outliers.

|  | All samples | | w/o *E*-*σ_max_* outliers | |
| --- | --- | --- | --- | --- |
| Group | *E* [MPa] | *σ_max_* [MPa] | *E* [MPa] | *σ_max_* [MPa] |
| Native | 59.21 ± 2.88 | 7.13 ± 0.31 | 58.83 ± 2.89 | 7.08 ± 0.31 |
| Triton X-100 | 4.42 ± 0.47 | 1.01 ± 0.09 | 123.51 ± 6.74 | 7.62 ± 0.47 |
| SDS | 83.24 ± 5.16 | 7.33 ± 0.44 | 80.10 ± 5.21 | 7.03 ± 0.45 |
| TnBP | 16.99 ± 1.67 | 1.18 ± 0.11 | 143.18 ± 6.87 | 8.14 ± 0.39 |
|  | w/o *E* outliers | | w/o *σ_max_* outliers | |
| Group | *E* [MPa] | *σ_max_* [MPa] | *E* [MPa] | *σ_max_* [MPa] |
| Native | 58.83 ± 2.89 | 7.08 ± 0.31 | 58.83 ± 2.89 | 7.08 ± 0.31 |
| Triton X-100 | 118.28 ± 6.14 | 7.89 ± 0.46 | 123.51 ± 6.74 | 7.62 ± 0.47 |
| SDS | 80.10 ± 5.21 | 7.03 ± 0.45 | 83.24 ± 5.16 | 7.33 ± 0.44 |
| TnBP | 143.18 ± 6.87 | 8.14 ± 0.39 | 143.18 ± 6.87 | 8.14 ± 0.39 |

**Table S8.** Average and standard deviation values of the maximum average strain deformations during the elastic stage within the four FL samples investigated through DIC analysis. e_yy_ corresponds to the longitudinal strain, i.e., parallel to the tensile direction and collagen fibers, e_xx_ corresponds to the transversal strain deformation, i.e., perpendicular to the tensile direction, e_VM_ refers to the Von Mises equivalent strain. For the longitudinal (e_yy_) and Von Mises (e_VM_) strains, we report maximum positive (tension) values, while for the transversal (e_xx_) strain we report minimum negative (compression) values. Both overall values as well as values is the three smaller sub-regions (R_0_, R_1_, R_2_; see Figure 7) are reported.

| Sample | Focus area | e_yy,max_ [-] | e_xx,min_ [-] | e_VM,max_ [-] |
| --- | --- | --- | --- | --- |
| Nat J4 | R_0_-Top | 0.0535 ± 0.0196 | -0.1502 ± 0.1067 | 0.1241 ± 0.0771 |
|  | R_1_-Center | 0.0221 ± 0.0108 | -0.1506 ± 0.0610 | 0.1106 ± 0.0639 |
|  | R_2_-Down | 0.0503 ± 0.0198 | -0.0255 ± 0.0375 | 0.0988 ± 0.0409 |
|  | Overall | 0.0357 ± 0.0198 | -0.0942 ± 0.0795 | 0.1111 ± 0.0640 |
| TRX F4 | R_0_-Top | 0.0334 ± 0.0026 | -0.0654 ± 0.0104 | 0.0618 ± 0.0038 |
|  | R_1_-Center | 0.0341 ± 0.0034 | -0.0942 ± 0.0366 | 0.0785 ± 0.0220 |
|  | R_2_-Down | 0.0397 ± 0.0072 | -0.0288 ± 0.0560 | 0.0728 ± 0.0262 |
|  | Overall | 0.0371 ± 0.0127 | -0.0422 ± 0.0541 | 0.0726 ± 0.0201 |
| SDS F3 | R_0_-Top | 0.1038 ± 0.0289 | -0.1803 ± 0.0264 | 0.3326 ± 0.0340 |
|  | R_1_-Center | 0.1352 ± 0.0526 | -0.1865 ± 0.0177 | 0.4208 ± 0.0436 |
|  | R_2_-Down | 0.1514 ± 0.0486 | -0.1875 ± 0.0358 | 0.4545 ± 0.0331 |
|  | Overall | 0.1235 ± 0.0731 | -0.1780 ± 0.0688 | 0.3581 ± 0.0899 |
| TnBP F3 | R_0_-Top | 0.0460 ± 0.0125 | -0.0162 ± 0.0034 | 0.0343 ± 0.0020 |
|  | R_1_-Center | 0.0590 ± 0.0070 | -0.0192 ± 0.0050 | 0.0564 ± 0.0110 |
|  | R_2_-Down | 0.0730 ± 0.0047 | -0.0488 ± 0.0089 | 0.0601 ± 0.0058 |
|  | Overall | 0.0710 ± 0.0186 | -0.0184 ± 0.0179 | 0.1086 ± 0.0997 |

**Table S9.** Summary of longitudinal and transversal elongations, as well as Poisson’s ratio ν, evaluated in correspondence of the maximum value of longitudinal elongation. E_l_ represents the elongation (ΔL/L) in the longitudinal direction, while E_0_ the elongation in the transversal direction. The Poisson’s ratio is obtained as the ratio (with opposite sign) between E_0_ and E_1_.

| Sample | E_1_ [-] | E_0_ [-] | ν [-] |
| --- | --- | --- | --- |
| Nat J4 | 0.1271 | -0.0295 | 0.232 |
| TRX F4 | 0.0834 | -0.0304 | 0.365 |
| SDS F3 | 0.2347 | -0.1177 | 0.501 |
| TnBP F3 | 0.1866 | -0.0375 | 0.201 |

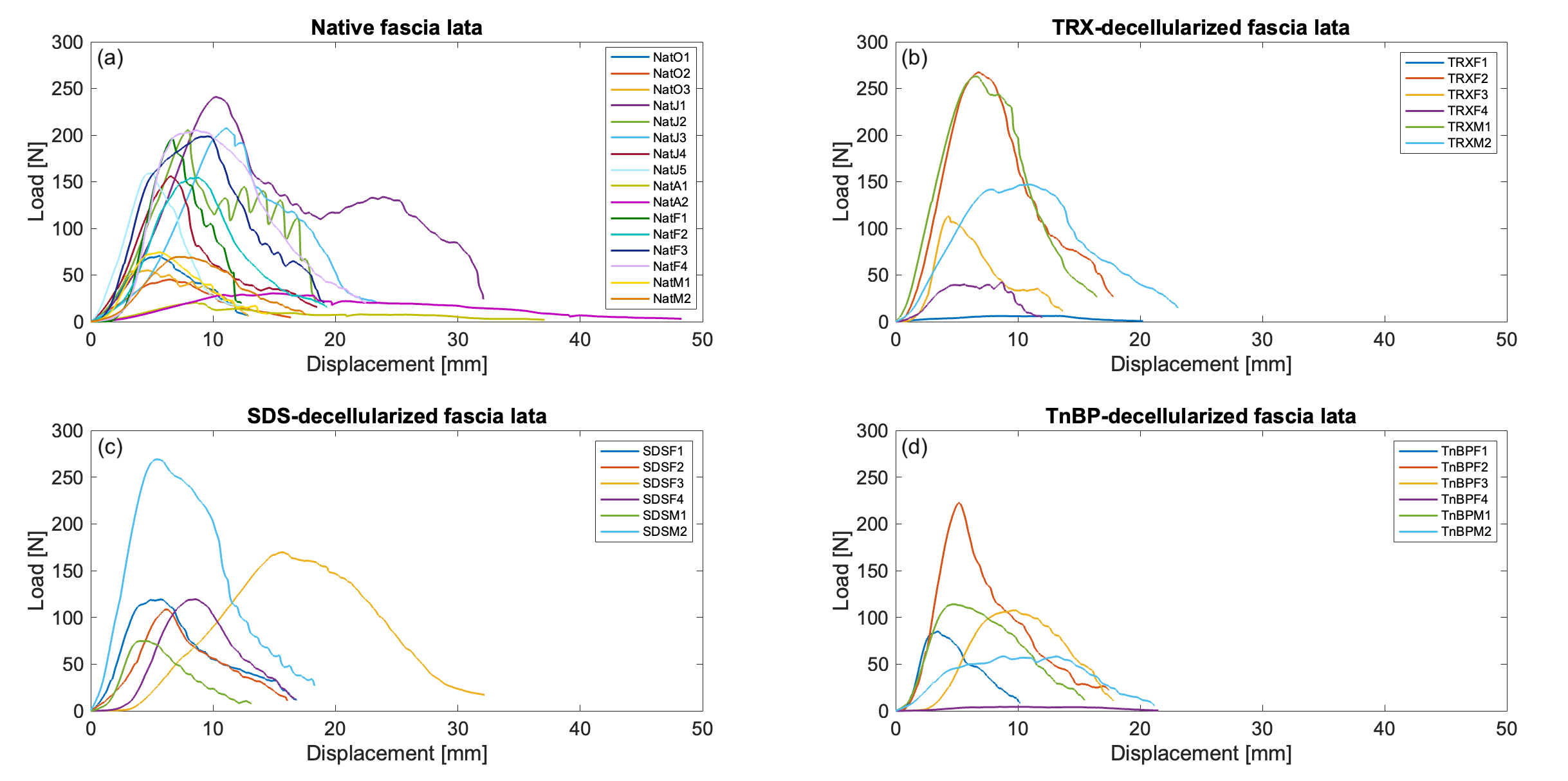

**Figure S1.** Full force-displacement curves for all FL specimens: (a) native samples; (b) samples treated with Triton X-100; (c) samples treated with SDS; (d) samples treated with TnBP. The colors of the curves refer to the different samples investigated during the various experimental campaigns.

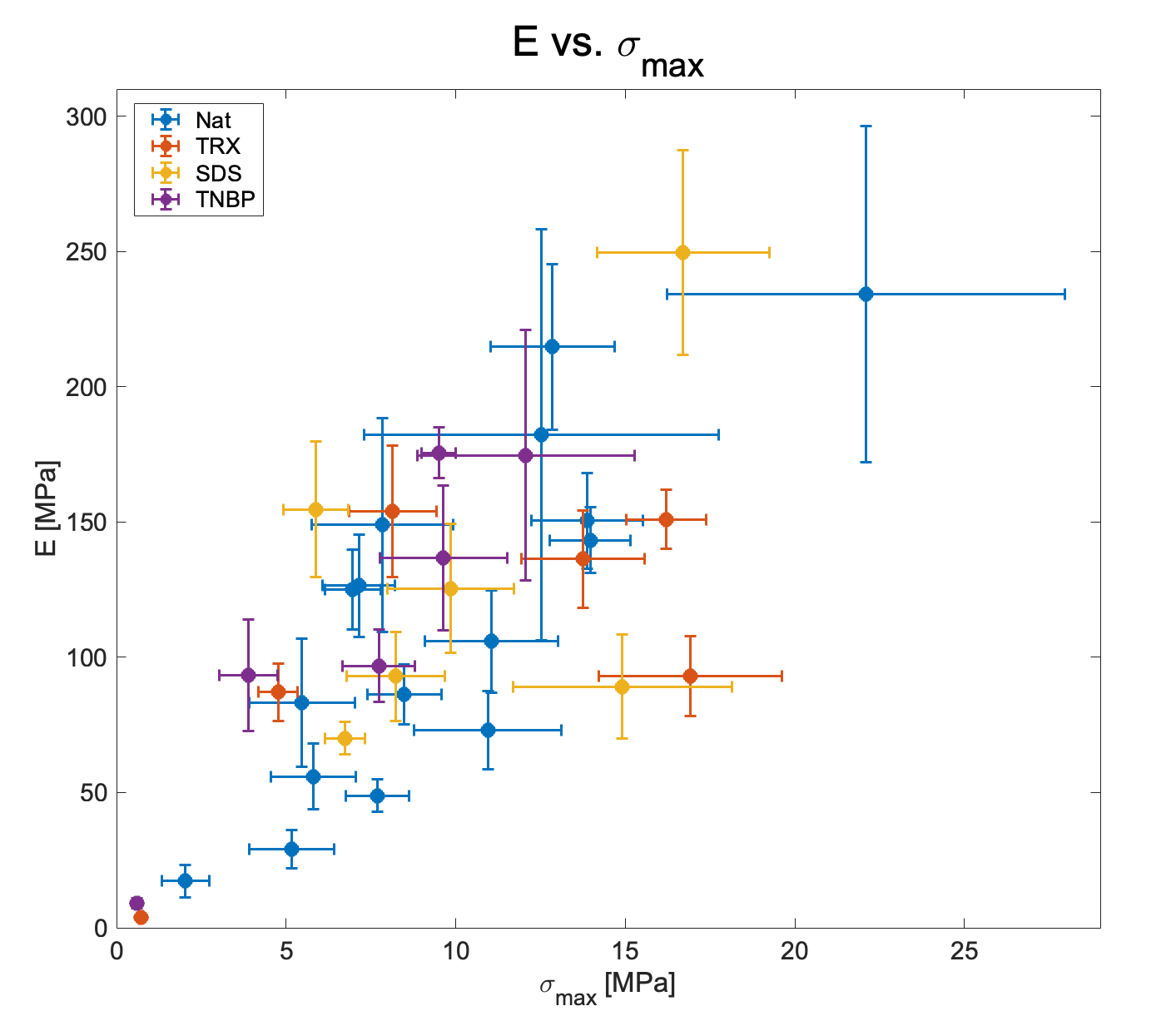

**Figure S2.** Correlation between elastic moduli E and ultimate stress σ_max_ for all FL samples (native in blue, treated with Triton X-100 in red, treated with SDS in yellow, treated with TnBP in purple). Each point represents average values of E and σ_max_ for each sample, with horizontal and vertical bars reflecting the amount of uncertainty. The Pearson correlation coefficient (PCC) between all E-σ_max_ average values is 0.724. Considering treatment groups separately, the PCC is 0.816 (native), 0.672 (Triton X-100), 0.521 (SDS), 0.935 (TnBP).

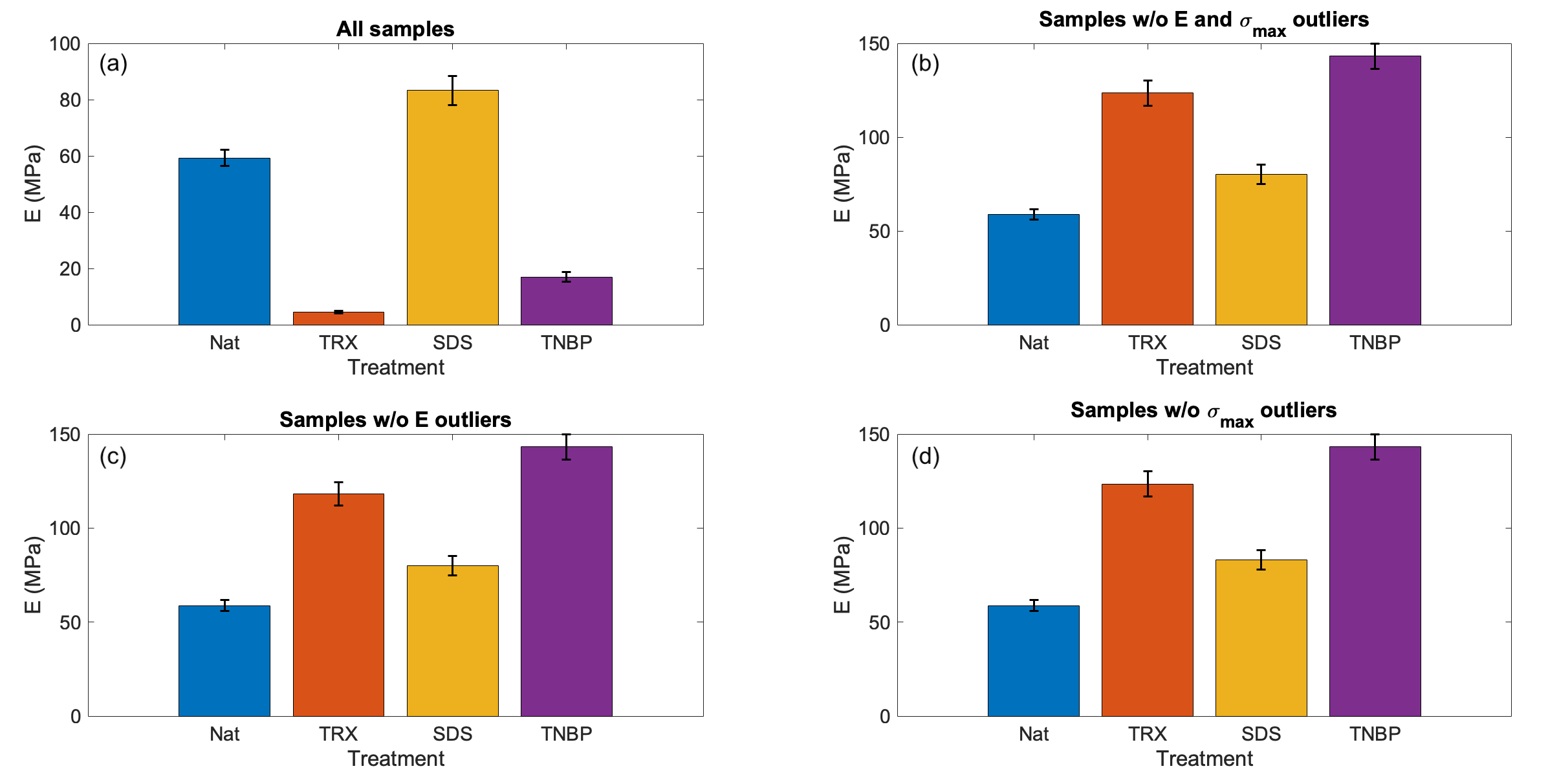

**Figure S3.** Values of elastic modulus E averaged over each FL treatment group (native in blue, Triton X-100 in red, SDS in yellow, TnBP in purple). Panel (a) reports average values and error bars of elastic moduli computed according to Eq. (S7b) for all 34 FL samples. The remaining panels show average values and error bars of elastic moduli not considering outliers (5^th^ and 95^th^ percentiles): (b) in both σ_max_ and E distributions; (c) only considering E values; (d) only considering σ_max_ values.

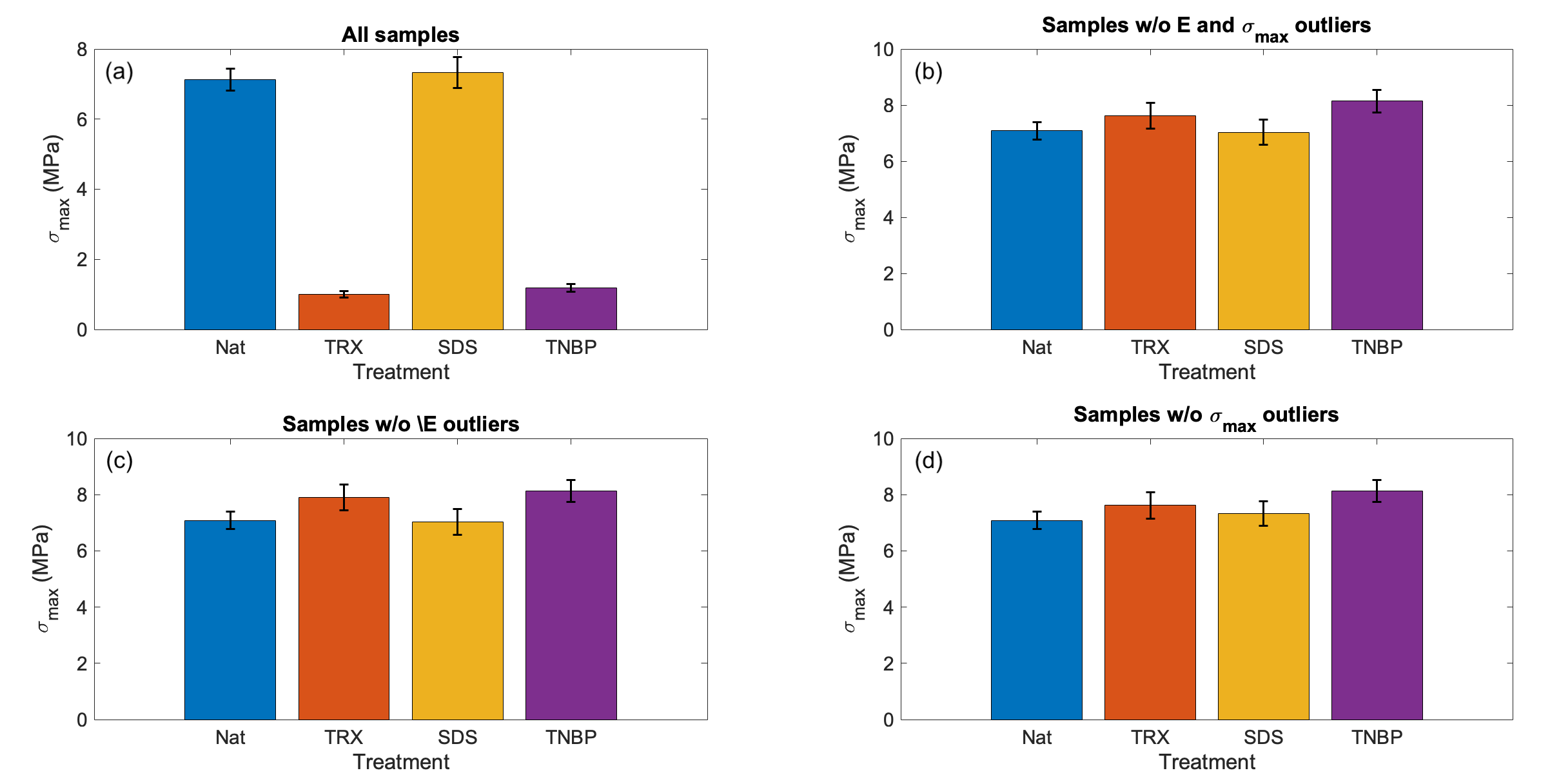

**Figure S4.** Values of maximum stress σ_max_ averaged over each FL treatment group (native in blue, Triton X-100 in red, SDS in yellow, TnBP in purple). Panel (a) reports average values and error bars of maximum stresses computed according to Eq. (S7a) for all 34 FL samples. The remaining panels show average values and error bars of maximum stresses not considering outliers (5^th^ and 95^th^ percentiles): (b) in both σ_max_ and E distributions; (c) only considering E values; (d) only considering σ_max_ values.
